# Resource Mismatch in Phototrophic Subaerial Biofilms and Resulting Phototroph/Heterotroph Ratios

**DOI:** 10.64898/2026.10.16.759371

**Authors:** Alberto Tenore, Fabiana Russo, Judy Jacob, Jean-David Grattepanche, Isaac Klapper

## Abstract

Photosynthesis-driven microbial communities are frequently observed on air-exposed stone such as, for example, marble surfaces. These environments are stark, subject to varying water, temperature, UV and other stresses. As a consequence, resident microbes likely have significant maintenance requirements. Those microbes however, have access to plentiful energy when conditions are favorable for photosynthesis. High rates of photosynthesis for purposes of energy generation may also generate excess organic carbon which can be released as extracellular overflow, becoming available for heterotrophic organisms. Here we propose a simple model to explore possible consequences of this hypothesized resource mismatch. The model predicts that heterotrophs can invade purely phototrophic communities, with the resulting mixed community eventually approaching a phototroph/heterotroph ratio at steady state as a consequence of stoichiometric balance. Results from field observations at three geographically separated sites (Lincoln Memorial, Washington D.C., Merchants’ Exchange, Philadelphia PA, and the Federal Hall National Memorial, New York NY), analyzed using two independent methods, are seen to match model predictions, demonstrating both cyanobacterial dominated communities as well as, mostly, mixed communities with a heterotrophs-to-cyanobacteria biovolume ratio of approximately 0.4.

## 1 Introduction

Air-exposed surfaces often host microbial communities, e.g., Figure 1, called subaerial biofilms (SABs), even under seemingly difficult conditions for life [25]. Outdoor surfaces in particular are often exposed to extremes in temperature as well as poor nutrient and water conditions. Extremophilic cyanobacteria, however, are capable of colonizing such environments, using sunlight and CO_2_ to fix carbon and produce energy, at least when sufficient water is available. Notably, though, despite their apparent self-sufficiency, bulk analyses of phototrophic (i.e., based on light for energy and carbon fixation) SABs generally reveal multispecies communities, where photosynthesizers are observed in combination with various types of heterotrophic microorganisms that are able to utilize photosynthetically produced organic carbon sources [7, 8, 13, 15].

**Figure (1).**
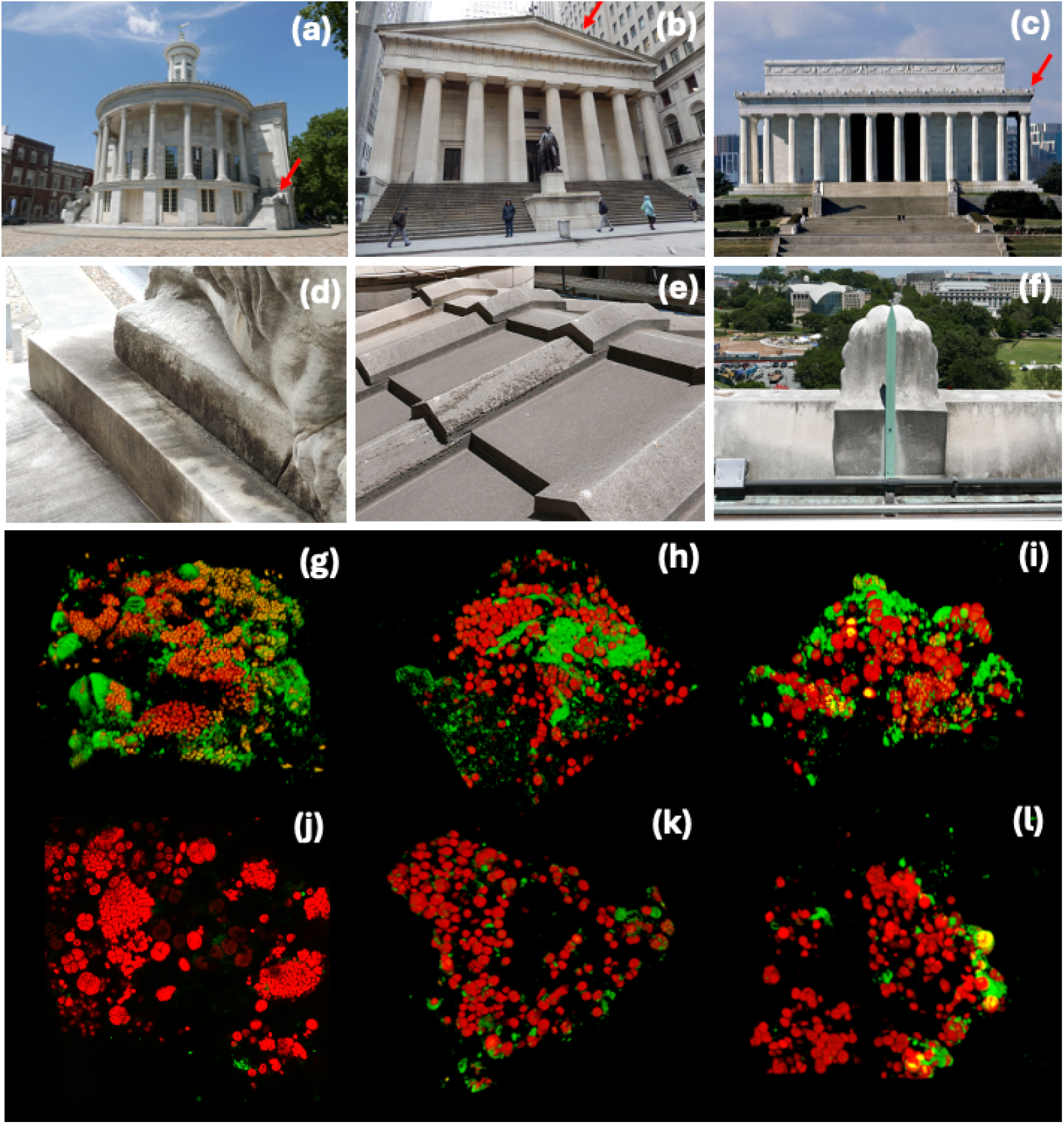
Confocal microscopy images of SAB samples from the Merchants’ Exchange Building, Philadelphia (a) and (d), Federal Hall National Memorial, New York City (b) and (e), and the Lincoln Memorial (c) and (f). (Top row) photos of the three sites, with red arrows pointing to sampling areas. (Middle row) close-ups of corresponding sites, with grayish areas containing extensive SABs. (Bottom) laser confocal microscopy images from samples. Cyanobacteria exhibit red autofluoresence from their photosynthetic machinery. SYTO 9 produces green fluorescence associated with DNA in heterotrophic cells and extracellular DNA. Images (g), (h), (i), show “typical” mixed cyanobacteria-heterotroph communities from the Merchants’ Exchange Building, Federal Hall National Memorial, and the Lincoln Memorial 3 respectively. Images (j), (k), (l), show rarer, mostly cyanobacteria communities, also from the Merchants’ Exchange Building, Federal Hall National Memorial, and the Lincoln Memorial respectively.x

While SAB heterotrophs require a reliable organic carbon source, e.g., cyanobacterial byproducts in the case of light-exposed SABs, to survive more than transiently, conversely there is also some evidence that cyanobacteria can benefit from heterotrophic activity. Such benefits are known to occur in aqueous environments in various ways [16]. Note that SABs, unlike typical aqueous communities, have the property that nonvolatile compounds are liable to accumulate because of the difficulty of transporting them away, suggesting a helpful role for heterotrophs in preventing byproduct inhibition. One possibility, for example, is that acidic cyanobacterial organic byproducts [4, 24], like glycolic acids, may inhibit cyanobacterial-only SABs; hence an advantage of heterotrophic partners.

SABs are, often, slow-growing and tightly acclimated to their environments, and hence are challenging systems to study either in the laboratory or in the field. As such, they are attractive targets for models. In fact, their extreme environmental circumstances may have the consequence that simple, physically based modeling assumptions can be effective. In this spirit, then, we present here a simple model, informed by field observations, of two phototrophic SAB communities. The model combines diel water and light conditions (both are only available, typically, intermittently over a 24 hour cycle) with longer time scale cyanobacteria-heterotroph interactions via a supposed acidic organic byproduct of cyanobacteria. We expect, based on observations (e.g., Figure 2), that this second time scale is months or longer. Thus there is a separation of scales between diel and growth times (note: additional time scales, e.g., seasonal or longer, are possible but not considered here), which we exploit in the model analysis, and which also suggests further measurement targets.

**Figure (2).**
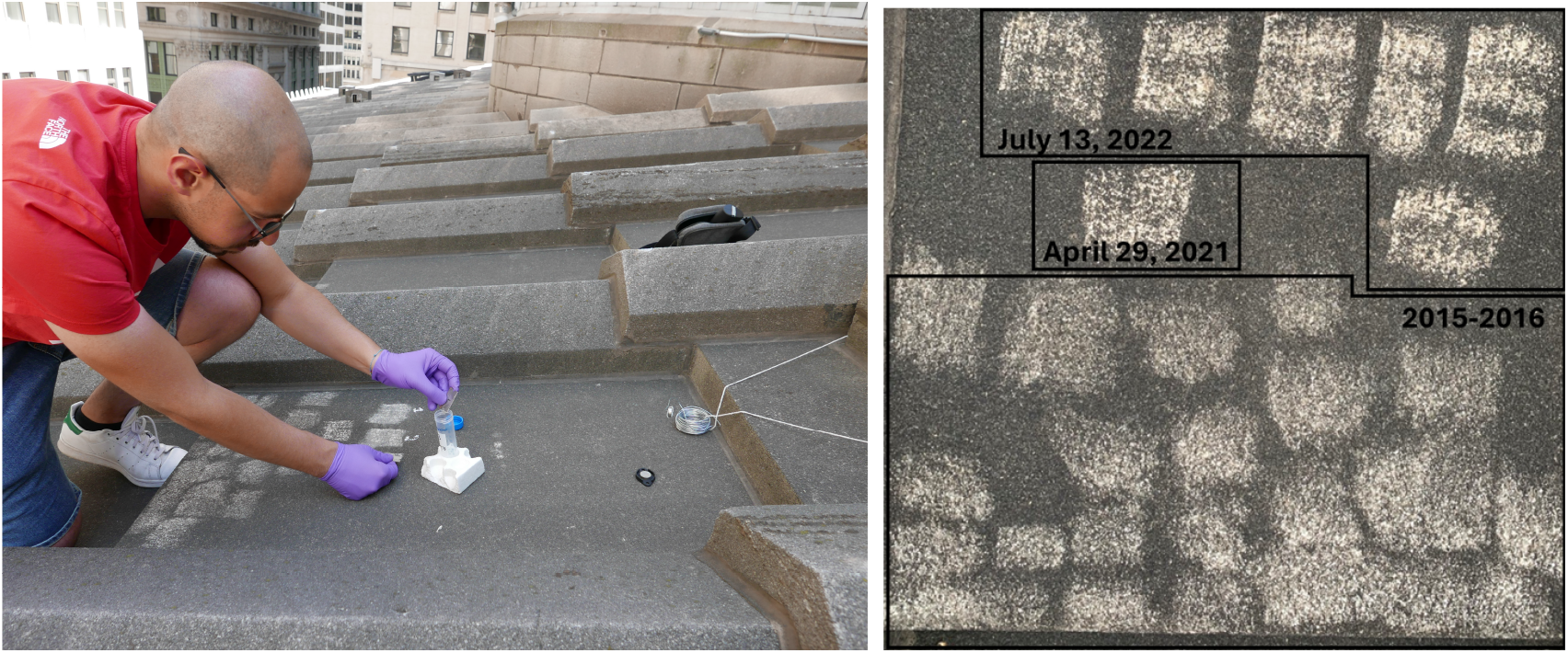
Sampling site on the roof of FH. (Left) A. Tenore (manuscript coauthor) collecting a scraping sample. White regions are newly scraped. Photo taken July 13, 2022. (Right) Photograph suggesting a long time scale for SAB regrowth. Sampling dates are indicated. Photo taken January 5, 2023.

**Figure (3).**
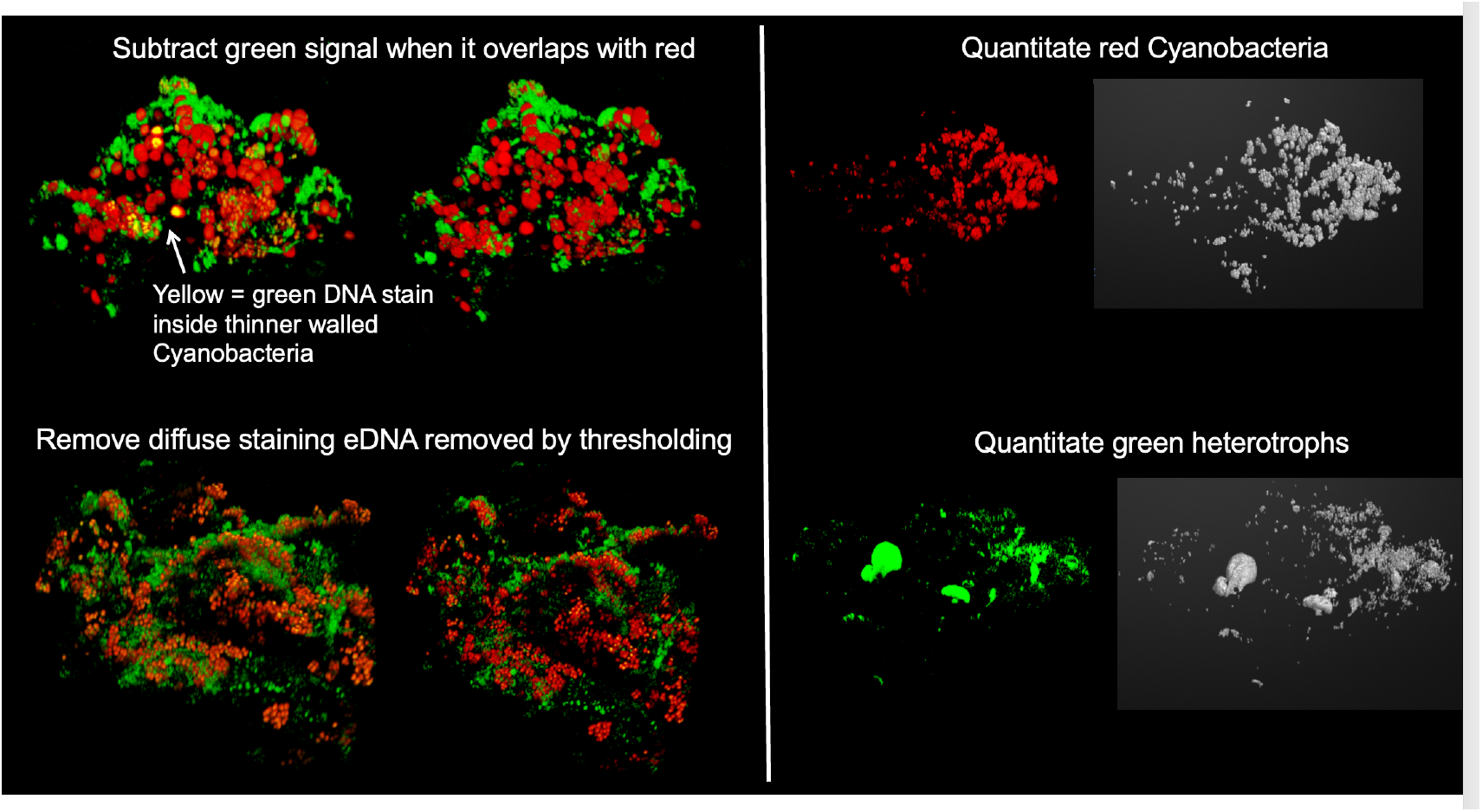
Biovolume estimation. Images contained four total signal types: red signal from photosynthetic machinery of cyanobacterial cells, yellow signal overlap of red and green DNA signal in cyanobacterial cells, small bright puncta of tightly packaged DNA inside small (*<* 3 *µ*m^3^) heterotrophs and diffuse signal from eDNA surrounding bacterial cells. (a) Subtraction of cyanobacterial green nucleus signal transitioning from yellow (green and red overlap) to red signal only. (b) Removal of eDNA from heterotroph signal. (c) Example of cyanobacterial and (d) heterotroph volumes detected by the Matlab connections algorithm-based method. Note different representative biofilms with easily distinguished details were chosen for (a) (b) and (c/d).

## 2 Field Observations

We present results based on samples of phototrophic SAB communities collected from marble at Federal Hall National Memorial (FH), New York City, the Merchants’ Exchange (ME), Philadelphia, and the Lincoln Memorial (LM), Washington, D.C., see Figures 1 and 2. These are all urban sites administered by the National Park Service: construction of FH was completed in 1842 using marble from Westchester County, New York, ME was completed in 1834 using marble from Montgomery County, Pennsylvania, and LM was completed in 1922 primarily using Colorado Yule marble for its exterior. FH is approximately 130 km from ME, which in turn is approximately 200 km from LM.

### 2.1 Methods

Methods are briefly outlined here; see references [2, 9, 23] for more detailed descriptions.

#### SAB Sampling

SABs on marble stone surfaces were collected from FH. ME, and LM by gently scraping an approximately 2×3 cm^2^ area of the surface patina with a sterile razor blade. The resulting granular material was used for microscopy and sequencing. For ME, samples analyzed in this study were collected from the base of the north lion statue. Sampling included one area in June 2021, three areas in July 2021, two areas in August 2021, and two areas in February 2022 (eight areas total). For FH, samples were collected from three areas in April 2022, six areas in July 2022, and four areas in March 2024 (13 areas total). For LM, samples were collected in October 2024. Individual samples were used only for microscopy or for sequencing, but not both.

#### Confocal microscopy

Samples intended for microscopy were immediately placed in dry ice and then later stored at *−*80 ^*o*^C. Samples were imaged by laser scanning confocal microscopy. Particles were stained with SYTO 9 to visualize DNA. SYTO 9 produces strong green fluorescence in heterotrophic cells but substantially weaker signal in cyanobacteria, with detectable green fluorescence in only a subset of cyanobacterial cells. SYTO 9 also stained extracellular DNA (eDNA) surrounding heterotrophic communities. Cyanobacteria were visualized by red autofluorescence of photosynthetic machinery. Imaging was done using a Leica SP5 microscope equipped with a TCS confocal system using a 63× objective at 1024×1024-pixel resolution. To quantify heterotrophic and cyanobacterial biomass, red- and green-channel thresholds were determined by visual adjustment. The green channel was subsequently processed to isolate the heterotrophic signal by removing green fluorescence associated with cyanobacterial cells and extracellular eDNA. Cyanobacterial and heterotrophic biovolumes were then quantified using a modified MAT-LAB connected-components algorithm [18]; code and additional methodological details are available at [17].

#### Sequencing

In the case of samples used for sequencing, RNA was extracted and reverse transcribed into cDNA to characterize the active microbial communities. The V4-V5 region of the bacterial 16S rRNA gene was amplified and sequenced using the Illumina MiSeq platform. Sequence data were processed through a custom Python pipeline for quality filtering, OTU clustering, and taxonomic assignment. OTUs were classified as phototrophic or heterotrophic based on their phylogeny [9]. For each sample, OTU relative abundances were first aggregated at the bacterial order level. Each order was then assigned to one ecological category based on its predominant ecological characteristics. The relative abundances of all orders within a category were summed, and these sums determined the size of the corresponding segments in stacked bar plots.

### 2.2 Results

Scraping samples were taken from three sites to characterize SAB community composition, specifically relative biovolumes of cyanobacteria and heterotroph inhabitants. See Figures 4 and 5 for imaging and sequencing results, respectively. Figure 4, based on processing of scanning confocal microscopy images, shows ratios of heterotroph to cyanobacteria biovolume counts over individual SAB colonies from the three sampled sites. Most SAB colonies have, approximately, a ratio of 0.32-0.42 (approximately 70-75% cyanobacteria, 25-30% heterotrophs), except for a small subset at ratio approximately 0.1 (approximately 90% cyanobacteria, 10% heterotrophs).

**Figure (4).**
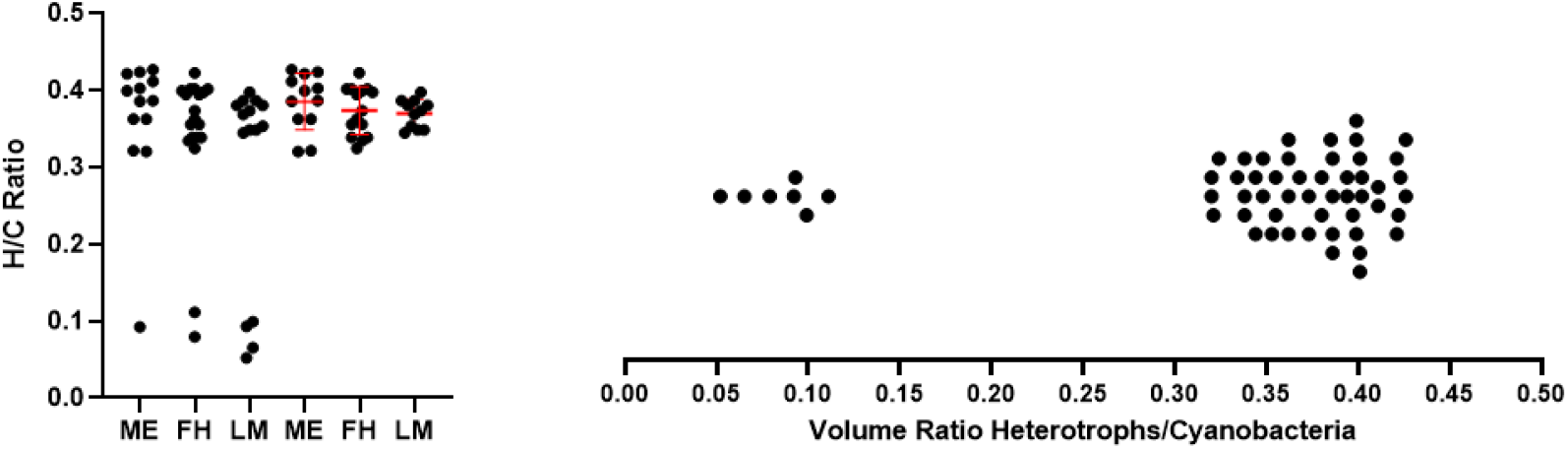
(Left) ratio of heterotroph biovolume to cyanobacterial biovolume measured from samples collected from ME, FH, amd LM; each dot corresponds to one sample. Dots above the first three ticks include all measured samples; dots above the next three exclude the small ratio outliers, but include horizontal bars indicating means and standard deviation ranges (excluding outliers). (Right) data from all three sites combined, with each dot corresponding to a sample with heterotroph/cyanobacteria volume fraction indicated by the value of the horizontal axis underneath.

**Figure (5).**
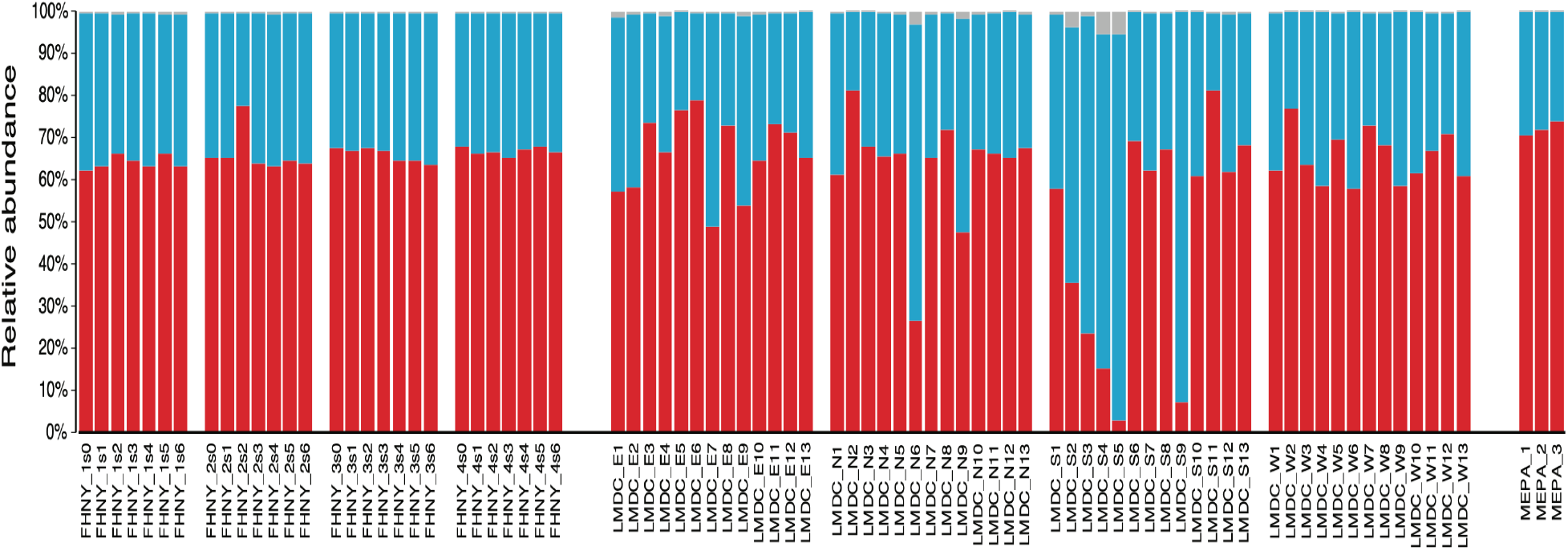
Functional composition of bacterial communities across three marble monuments. Stacked bars show relative abundances (% of total reads) of Phototrophs (red), Heterotrophs (blue), and Others (grey; including chemolithotrophs, predators, intracellular bacteria, and taxa of unknown ecology). Functional groups were assigned based on the typical metabolism and ecology of each bacterial order; see [9] for more detail. Samples are from FH (FHNY; digits 1–4 indicate roof sampling locations), LM (LMDC; E, N, S, and W indicate roof orientation), and ME (MEPA). Some samples, particularly from the sunnier south side of LM, showed distinct compositions, the causes of which are unresolved.

Microscopy-based observations are supported independently (both in terms of the methodology, and in terms of samples – separate samples are used for imaging vs sequencing) by ratios of heterotroph to cyanobacterial 16S cDNA-based reads, see Figure 5, with median values and ranges reported in Table 1. Note that, because of the processing method, each bar graph in Figure 5 is in fact already an average over many individual SAB colonies of the sort counted as individual points in Figure 4. Also note that the two methods are measuring somewhat different quantities, so we can’t rule out some level of coincidence in the similar ratios, but the fact that both methods show, mostly, a constant ratio is unmistakable and striking.

**Table (1).** Median relative abundances (% of total reads), with ranges in brackets, for the stacked bar graphs shown in Figure 5.

|  | Federal Hall | Lincoln Memorial | Merchants' Exchange |
| --- | --- | --- | --- |
| **Phototrophs** | 65.38% [62.19-77.64%] | 65.4% [2.79-81.29%] | 72.02% [20.72-73.90%] |
| **Heterotrophs** | 34.10% [22.10-37.30%] | 34.30% [18.51-92.55%] | 27.85% [25.87-29.07%] |
| **Others** | 0.49% [0.26-0.69%] | 0.35% [0.03-5.35%] | 0.20% [0.13-0.23%] |

While all samples are taken from marble building structures in the north-eastern United States, the three sites have some significant differences in local climate (in addition to likely differences in marble material characteristics). FH is located in a warm, urban canyon in lower Manhattan, while LM is located in a relatively open and wet environment, close to the Washington D.C. tidal basin; conditions at ME are likely somewhat intermediate. Note additionally that samples on LM roof are taken from each of its four sides (north, east, south, west) over which temperature and relative humidity conditions can vary considerably.

## 3 SAB Community Model

Observations suggest three SAB community types: (1) a null community, (2) a cyanobacteria-only (or almost only) community, and (3) a mixed cyanobacterial-heterotroph community. We construct a simple model, based on available evidence, to explore a possible succession among these three types, as follows. An uncolonized surface (type 1 community) can be susceptible to invasion of cyanobacteria (type 2 community) if environmental conditions are sufficiently favorable, with a quantification of environmental conditions made below. Note that heterotrophs depend on cyanobacterial byproduct for an organic carbon source, and so can’t persist without a sufficiently large cyanobacterial host community. However, as the cyanobacterial biomass increases, it may become possible for heterotrophs to invade (type 3 community).

The model tracks populations of cyanobacteria *C*(*t*) [molC m^−3^], heterotrophs *H*(*t*) [molC m^−3^], and a cyanobacterial byproduct/heterotroph nutrient *S*(*t*) [molC m^−3^]. Cyanobacteria grow and produce a byproduct (which also is supposed to be a cyanobacterial inhibitor) at rate proportional to growth which is used by heterotrophs as a nutrient, see Figure 6 for a schematic. In equation form, this amounts to

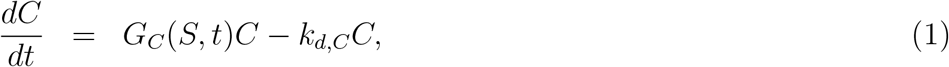

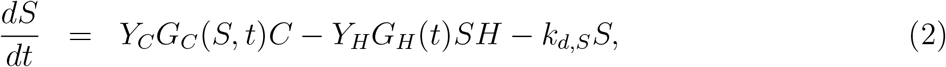

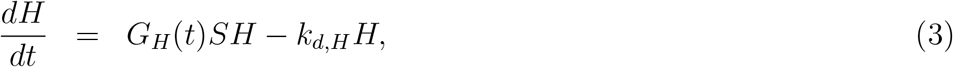

where *G*_*C*_(*S, t*) (a decreasing function of *S*) and *G*_*H*_ (*t*) are positive specific growth rates, to be described in detail below, *k*_*d,j*_ are positive decay rates, and *Y*_*j*_ are positive yield conversion coefficients. Note the explicit dependence of *G*_*C*_ and *G*_*H*_ on time, due to the diel cycle (and also in principle seasonal or other time dependencies). We suppose below that *G*_*C*_(*S, t*) and *G*_*H*_ (*t*) are periodic in *t*, with period of one day, neglecting other possible time variations. A more detailed characterization of community metabolic activity, which can depend on environmental conditions, can be included (see, e.g. [22, 20]) but is not necessary here.

**Figure (6).**
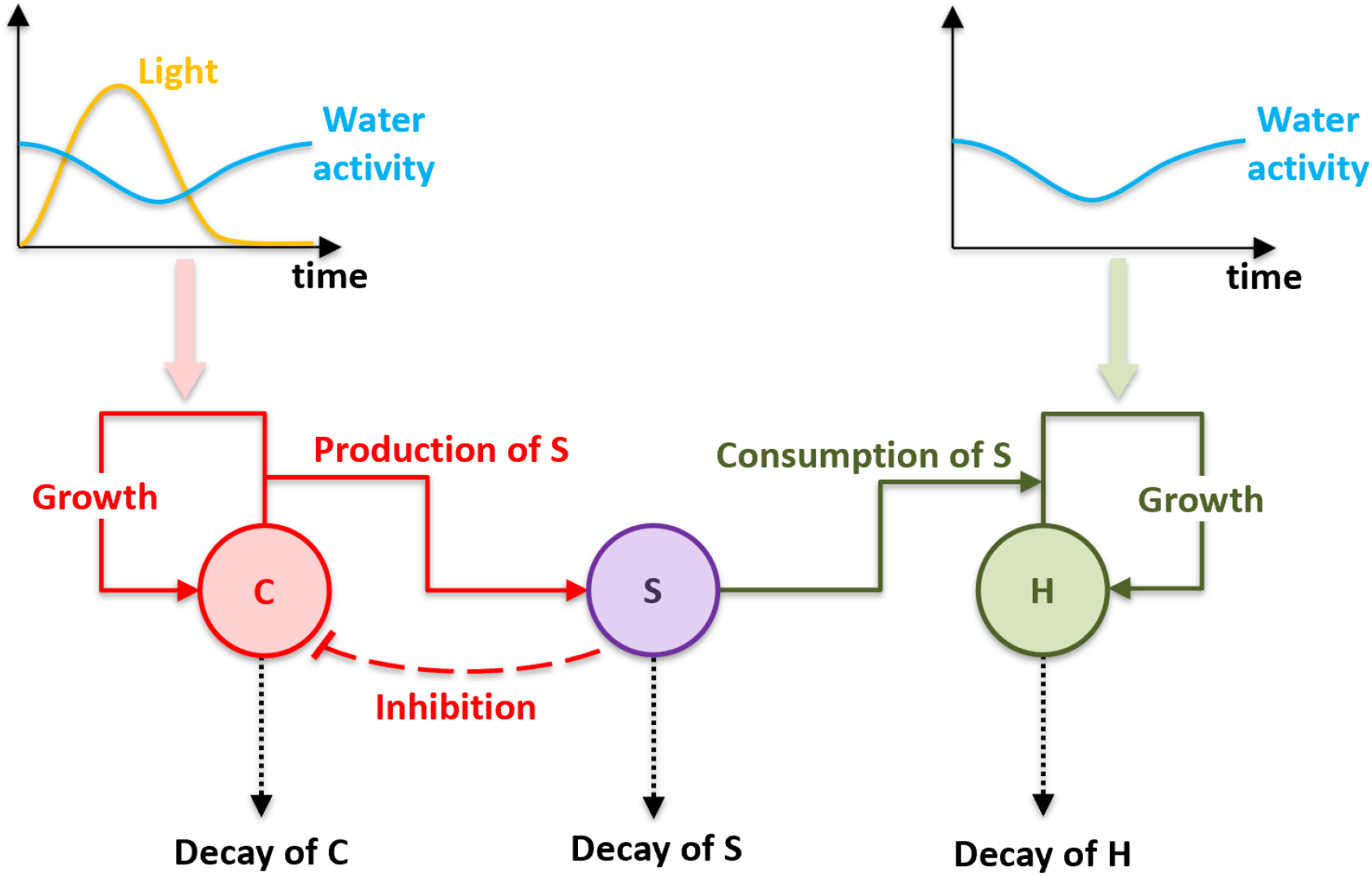
Schematic representation of the model community metabolic network showing the three components cyanobacteria (C), byproduct/nutrient (S), and heterotrophs (H) and their connections. A single diel cycle of light intensity (for cyanobacteria) and water activity (see Section 3.2) are shown at the top.

### 3.1 Stability of the Null Community

Equations (1)-(3) have a null steady state *C*_*N*_ = *S*_*N*_ = *H*_*N*_ = 0, i.e., the null community. Perturbing from this state, with small initial conditions 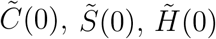, then we obtain

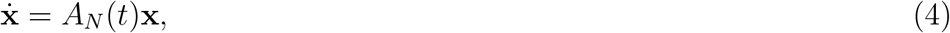

where **x** is the vector with components 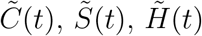 and

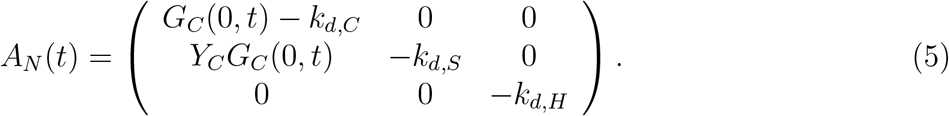

Note *A*_*N*_ (*t*) is P-periodic, where P is one day in whatever time units are being applied.

System (5) can be analyzed using ideas from Floquet theory [10], which states that, for a fundamental matrix solution *M* (*t*) of (4), there is a constant matrix Ω such that *M* (*t* + *P*) = *M* (*t*)Ω for any *t*. Observing then that *M* (*t* + *nP*) = *M* (*t*)Ω^*n*^, we see that if any eigenvalue of Ω has absolute value greater than 1, then *M* (*t*), and thus the non-viable steady state, exhibits instability; conversely, if all eigenvalues have absolute value less than 1, then the non-viable steady state is stable. (Matrix Ω depends on choice of fundamental matrix solution but its eigenvalues do not.)

Note that Ω = *M* ^−1^(0)*M* (*P*) = *M* (*P*) if *M* (0) = *I*. Denoting, then, *M* (*t*) as a fundamental matrix solution with *M* (0) = *I*, we find after integrating (4) over one period *P* that

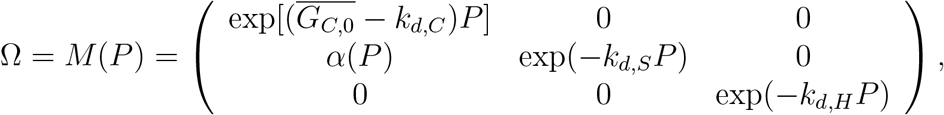

Where

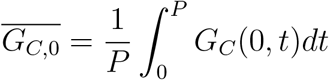

is the day-averaged uninhibited growth rate, and

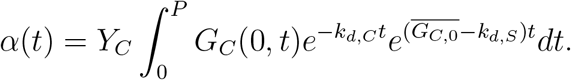

The eigenvalues 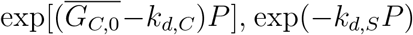, exp(*−k*_*d,S*_*P*), and exp(*−k*_*d,H*_ *P*) of *M*(*P*) indicate that the steady state *C* = *S* = *H* = 0 is linearly unstable if 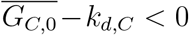 , and linearly stable if 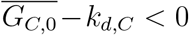. That is, a cyanobacterial population will be viable if its diel-averaged uninhibited growth rate 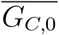 is larger than its decay rate *k*_*d,C*_ . (If the two rates are equal, then the cyanobacterial population cannot grow, as *G*_*C*_(*S, t*)*−k*_*d,C*_ *≤ G*_*C*_(0, *t*)*−k*_*d,C*_ for *S >* 0, so that 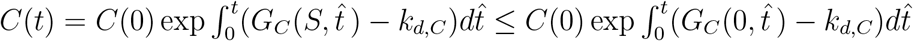,which is bounded in this case.) Note also that any solution with 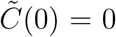 decays if *k*_*d,H*_ *>* 0, as does the 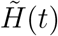 component of any solution at all. Thus the *C* = *S* = *H* = 0 solution is always linearly stable to invasion of heterotrophs.

### 3.2 Growth Rate Functions and Time Scaling

Details of the growth rates *G*_*C*_(*S, t*) and *G*_*H*_ (*t*) are in fact of some interest, as their t-dependence encodes extremes in daily environmental fluctuations and the S-dependence of *G*_*C*_ encodes an essential piece of the proposed ecology, namely byproduct inhibition. In particular, we focus on water cycling, with water conditions optimal during early morning because of condensation, see e.g. [23, 26]. Photosynthesis also depends on light availability. With these considerations in mind, we set

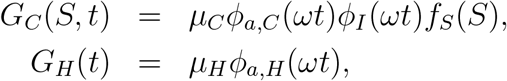

where *µ*_*C*_ and *µ*_*H*_ are maximum growth rates. Water limiters *ϕ*_*a,C*_ (*t*) and *ϕ*_*a,H*_ (*t*), both periodic with period 1 (the parameter *ω* sets the time scale), are functions with values in [0 1] that account for the effects of poor water conditions on growth of, respectively, cyanobacteria and heterotrophs. A value of 1 indicates good water conditions, with decreasing values for poorer conditions.

The water limiters are defined here as

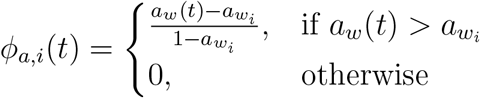

where *a*_*w*_(*t*) is the water activity, assumed to be periodic with period 1, and taking values in the range [0, 1], with *a*_*w*_(*t*) = 1 corresponding to ideal conditions. The factors 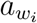are threshold values of water activity for cyanobacteria and heterotrophs, below which growth cannot occur.

Light limiter *ϕ*_*I*_(*t*), also periodic with period 1, is defined here by

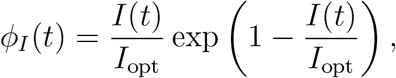

where *I*_opt_ denotes the optimal light intensity, and *I*(*t*), the light intensity in time, is supposed periodic with period 1, following [14]. The light limiter takes its maximum value 1 in ideal conditions, and allows the possibility of oversaturation, though that is not a significant issue here because water conditions are poor when light intensity is high. See Figure 7 for examples of *I*(*t*), *a*_*w*_(*t*), *ϕ*_*I*_(*t*) and *ϕ*_*a,C*_ (*t*), *ϕ*_*a,H*_ (*t*).

**Figure (7).**
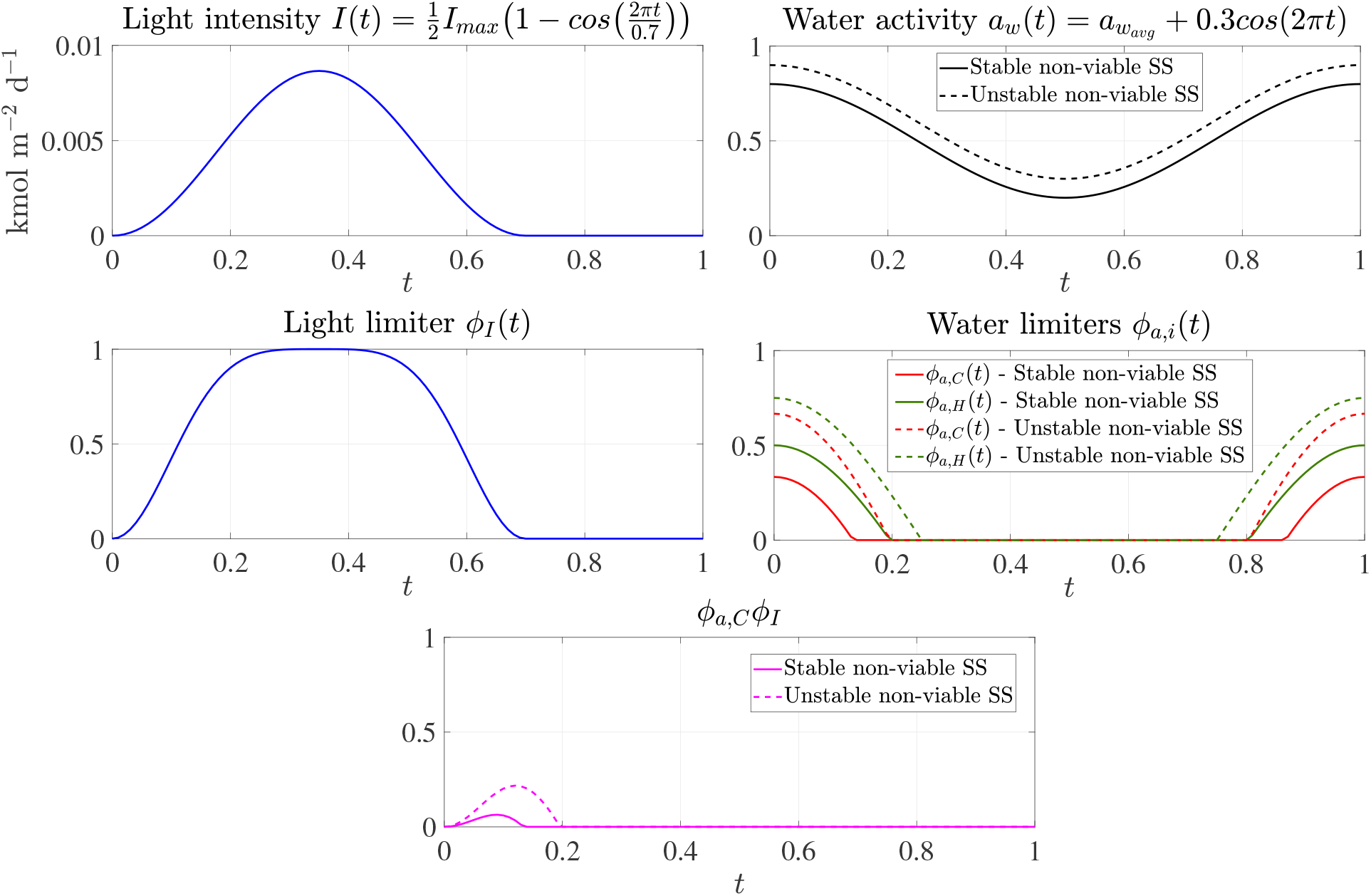
(Top) imposed light intensity and water activities as functions of time of day (*t* = 0 corresponds to daybreak), with *I*_*max*_ and 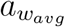 denoting the maximum light intensity and the average value of water activity, respectively. (Middle) light limitation and water activity limitation functions, computed from given intensity and activity functions. (Bottom) product of light and water activity limitation functions *ϕ*_*a,C*_ (*ωt*)*ϕ*_*I*_(*ωt*). Water activity in the early morning largely determines whether the non-viable steady state *C* = *S* = *H* = 0 is unstable.

The function *f*_*S*_(*S*) is a byproduct inhibition factor, which is supposed positive and monotone non-increasing, with *f*_*S*_(0) = 1 and *f*_*S*_(*S*) approaching zero as *S* increases. For computations, we use

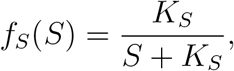

where *K*_*S*_ is an inhibition half-concentration. Byproduct inhibition is not a primary feature of the model, but is included to allow the presence of heterotrophs to improve conditions for cyanobacteria, as has been reported in [27].

We define the unitless averages

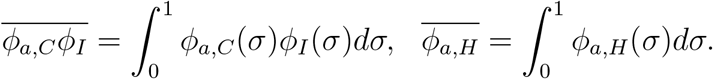

Setting the system growth time to 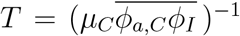, and scaling time in equations (1)-(3) by

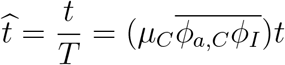

results in, after dropping hats,

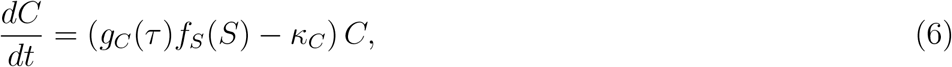

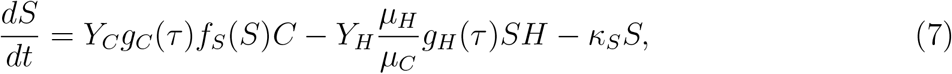

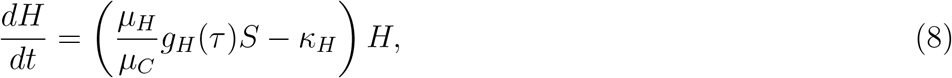

where *τ* = *t/ϵ* with 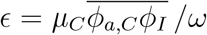, the ratio of the environmental period to the cyanobacterial population growth time. Parameters 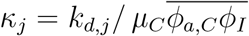 are ratios of cyanobacterial growth to decays times, and

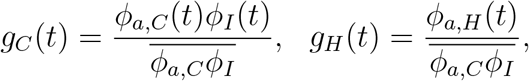

are scaled growth functions. Note that *g*_*C*_(*t*) has average 1. In what follows, we will suppose *ϵ* to be small, i.e., population growth time is long compared to the environmental cycle time (1 day), see, e.g., Figure 2.

## 4 Viable Solutions

### 4.1 Two Time-Scale Asymptotics

Equations (6)-(8) involve two separated time scales, the diel cycle time and the longer growth time. Their ratio is measured by the nondimensional number *ϵ*. The short diel cycle, though of interest because of coupling of water and light conditions over the course of a day, makes analysis of longer time behavior awkward. We thus employ multiple time scale asymptotics. Supposing *ϵ* to be small we designate *t* be “slow” time and let *τ* = *t/ϵ* to be “fast” time, and assume the two to be independent asymptotically as *ϵ→*0. Then, setting

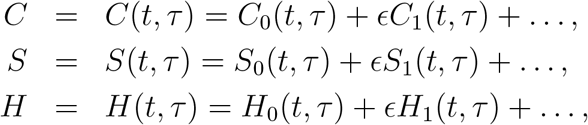

and noting

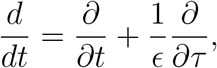

equations (6)-(8) become

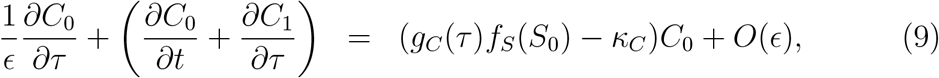

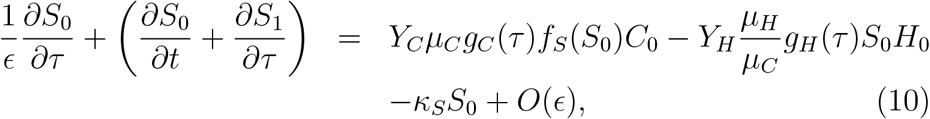

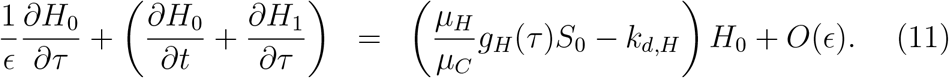

Note, as mentioned previously, dependencies *g*_*C*_ = *g*_*C*_(*τ, t*) and *g*_*H*_ = *g*_*H*_ (*τ, t*) are possible, i.e., slow environmental changes, e.g., seasonal or secular, are possible, though we will not consider such possibilities here.

To begin, we see at *O*(*ϵ*^−1^) that *∂C*_0_*/∂τ* = *∂S*_0_*/∂τ* = *∂H*_0_*/∂τ* = 0, i.e., *C*_0_ = *C*_0_(*t*), *S*_0_ = *S*_0_(*t*) and *H*_0_ = *H*_0_(*t*) are independent of fast time *τ* . At *O*(*ϵ*^0^),

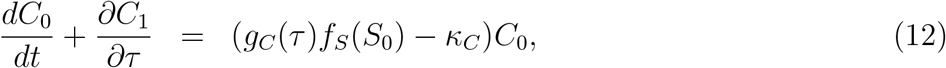

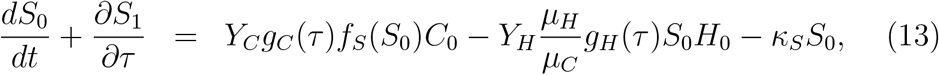

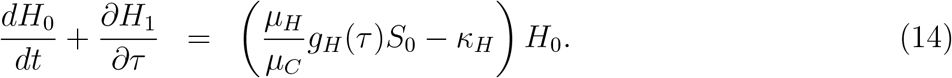

Integrating over fast time *τ* and letting *τ → ∞*, we obtain

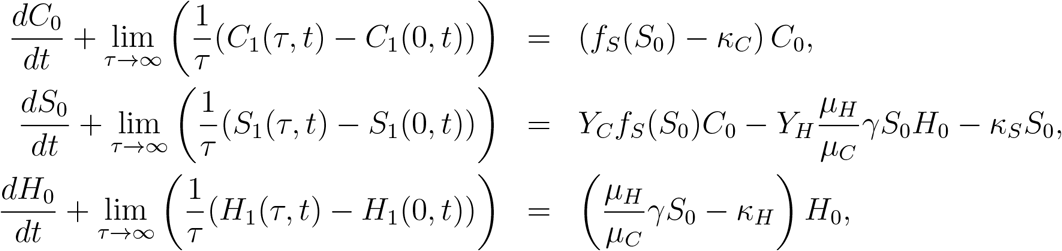

where 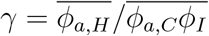. To avoid secular behavior in *τ* , we require the limits to be 0, thus obtaining equations

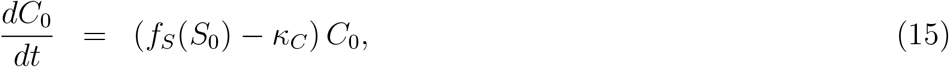

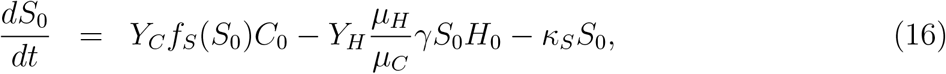

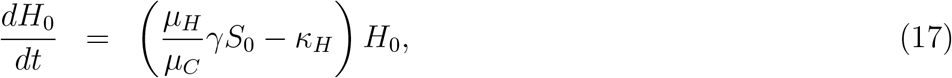

for *C*_0_(*t*), *S*_0_(*t*), and *H*_0_(*t*). See Figure 8 for a comparison of solutions to equations (6)-(8) with solutions of (15)-(17). Initial conditions *C*_0_(0) and *S*_0_(0) are set equal to the temporal mean values of *C*(*t*) and *S*(*t*) obtained over the first period of the corresponding simulation of equations (6)–(8), with *H*_0_(0) = 0. These choices ensure that the two models are initialized from comparable states while accounting for the periodic variability in (6)– (8).

**Figure (8).**
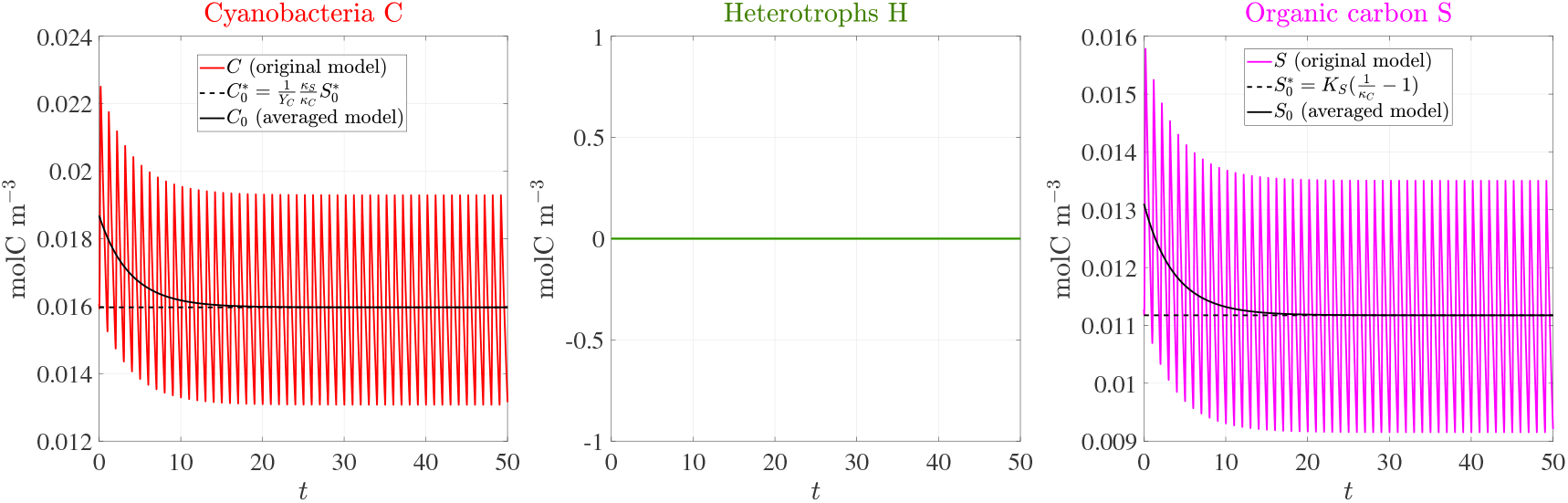
Comparison of solutions to equations (6)–(8) with solutions to the corresponding averaged equations (15)–(17).

### 4.2 Steady States of the Averaged Equations

Equations (15)-(17) have a non-viable solution *C*_0_ = *S*_0_ = *H*_0_ = 0. This state has the same stability characteristics as in the original model. In particular, in scaled quantities, a cyanobacterial population becomes viable, i.e., *C*_0_ = 0 is unstable, when *f*_*S*_(0) *−κ*_*C*_ = 1 *−κ*_*C*_ *>* 0 and not viable when *f*_*S*_(0) *−κ*_*C*_ = 1 *− κ*_*C*_ *<* 0; a heterotroph community is not viable whenever *κ*_*H*_ *>* 0.

There is also a second steady-state solution 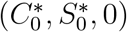, a monospecies cyanobacterial state, where 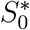 is determined by the relation

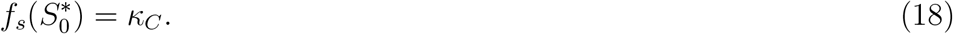

Since *f*_*S*_ is monotonically decreasing from 1 to 0, condition (18) will have exactly one solution 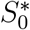, with 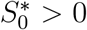, in the case that *κ*_*C*_ *<* 1, which is also the condition for instability of the *C*_0_ = *S*_0_ = 0 steady state. (If *κ*_*C*_ *>* 1 then (18) has no solutions, and there is no second steady state and the non-viable steady state is stable.) Given 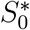, then

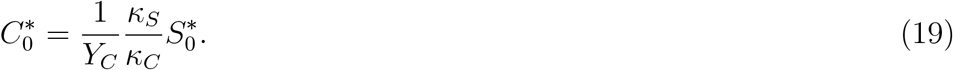

This monospecies steady state occurs when organic carbon *S*_0_ reaches a level so that growth is reduced by substrate inhibition to the point that it balances with decay. This steady state does not appear in the original equations (1)-(3) because that system is non-autonomous. Rather, a corresponding periodic solution is observed, at least numerically.

We also note that we don’t know, in actual phototrophic SABs, what limits SAB colony size. In particular, we don’t know that substrate inhibition would indeed do so, as it does in this model system. However, the existence of the monospecies cyanobacterial fixed point is not important to the main points of this paper, so can even be regarded as something of an artifact without impacting what follows in any significant way (it is in fact unstable in the most relevant case, see next paragraph). Conditions for its lack of stability to heterotroph introduction could still be of some secondary interest, though, as these conditions can suggest a delayed invasion of heterotrophs into an already established cyanobacterial SAB.

The second steady state, when it exists, has linearization matrix given by

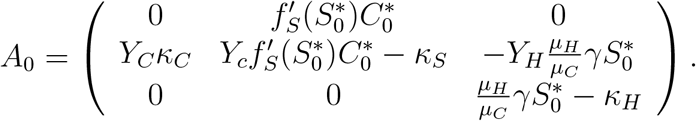

Note that perturbations in *C* and *S* only, i.e., those for which *H*_0_ = 0 is left unperturbed, decay as the *C*_0_-*S*_0_ plane is invariant under equations (15)-(17) and eigenvalues of the upper 2×2 submatrix of *A*_0_ (which coincide with two of the eigenvalues of *A*_0_) have negative real parts. The third eigenvalue of *A*_0_ is given by the lower right entry 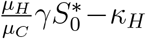, which may be positive or negative. We presume based on observations that there are heterotroph species for which this value is positive, through controlling *µ*_*H*_ or *γ* or *κ*_*H*_ , and so the monospecies steady state is generally unstable to invasion by heterotrophs. That is, the unstable parameter range is the biologically relevant one.

### 4.3 Multispecies Communities

Field observations show three types of communities: null-communities (bare rock, roughly), cyanobacterial-only communities (somewhat rarely), and mixed cyanobacterial-heterotrophic communities. We can see these three cases also in model dynamics, see Figure 9, in which a non-volatile community is linearly unstable to a cyanobacterial invasion which eventually, after organic byproduct accumulation, becomes susceptible to heterotroph invasion (i.e., a small positive perturbation of heterotrophs can grow). To further understand this progression, we study stability of a monospecies solution to equations (15)-(17) with *S*_0_(0) = *H*_0_(0) = 0 and *C*_0_(0) small, i.e., introduce a small cyanobacterial perturbation to the null state *C*_0_ = *S*_0_ = *H*_0_ = 0, with parameters such the instability condition 1*−κ*_*C*_ *>* 0 holds. (Bare rock itself can be susceptible to invasion where water conditions are sufficiently favorable, e.g., in the cracks between grains.)

**Figure (9).**
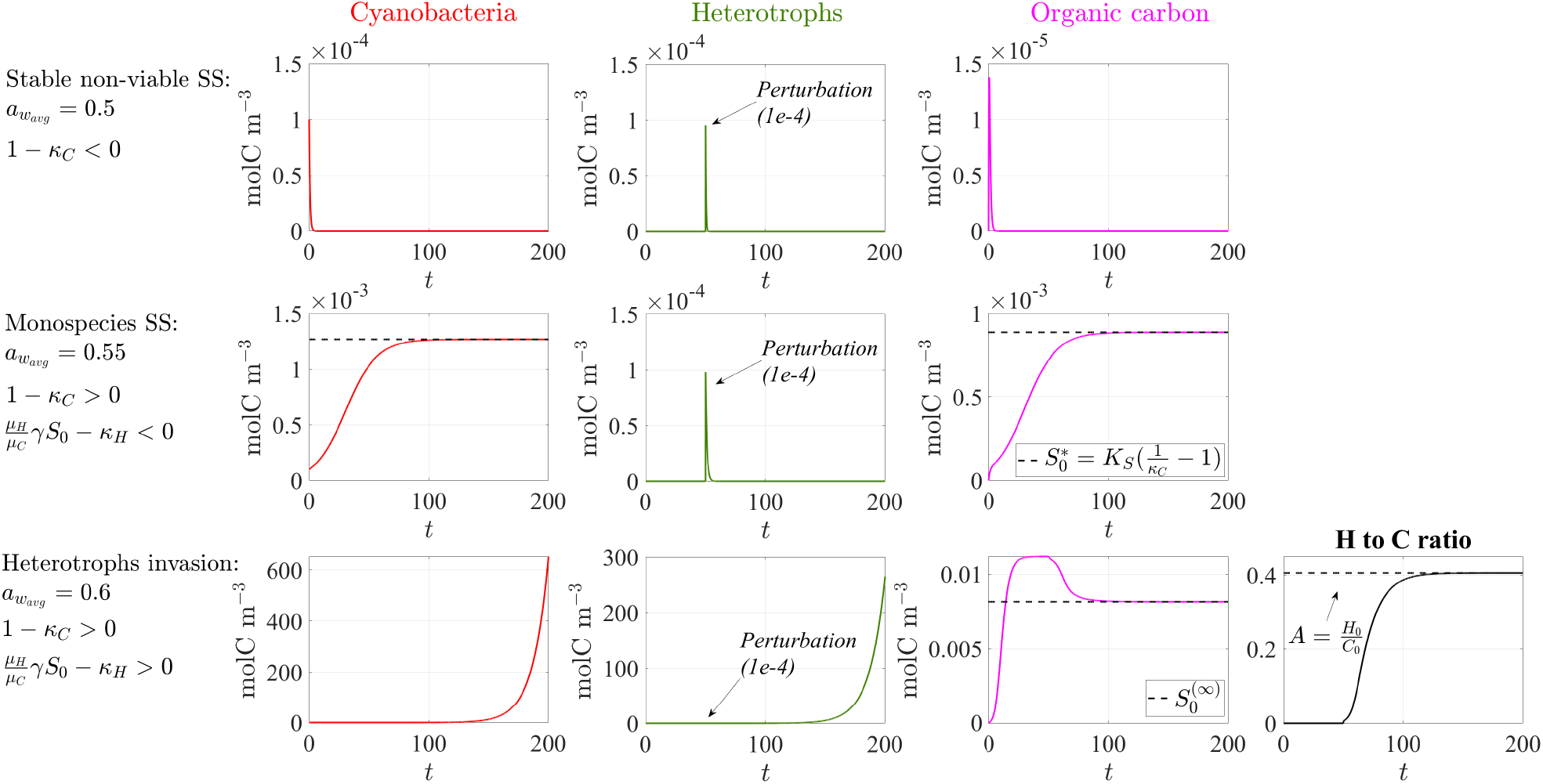
Numerical solution of equations (15)-(17), with cyanobacteria introduced at *t* = 0 d and heterotrophs introduced at *t* = 50 d. (Top) parameters are such that environmental conditions are non-viable. (Middle) parameters are such that environmental conditions are viable for cyanobacteria but non-viable for heterotrophs. (Bottom) parameters are such that environmental conditions are viable for both cyanobacteria and for heterotrophs. Note in this case that both cyanobacteria and heterotrophs grow but their ratio approaches a steady state.

Denoting the monospecies solution as (*C*_0_(*t*), *S*_0_(*t*), *H*_0_(*t*)) and setting

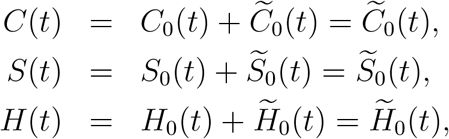

in the case *C*_0_ = *S*_0_ = *H*_0_ = 0, where quantities with tildes are considered small perturbations, we obtain the linearized equation

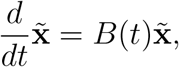

with

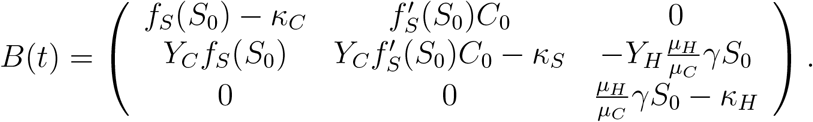

Note that 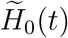 partially decouples; in particular

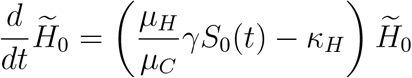

indicating growth when ((*µ*_*H*_ */µ*_*C*_)*γS*_0_(*t*) *− κ*_*H*_) *>* 0, which occurs for large enough *S*_0_(*t*). (Note: this quantity is monotonically increasing for early times; we will assume that parameters are such that it in fact becomes positive, because heterotroph invasion is generally observed.) Invasive species with *κ*_*H*_ */*(*γµ*_*H*_ */µ*_*C*_) small will be favored, i.e., with some kind of combination of high efficiency *µ*_*H*_ */κ*_*H*_ and a relatively low water activity tolerance number *γ* is desirable.

Once successful invasion occurs, we wish to characterize long time behavior by looking for solution (*C*_0_(*t*), *S*_0_(*t*), *H*_0_(*t*)) to approach a state with *H*_0_(*t*)*/C*_0_(*t*) = *A*, 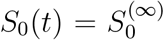, where *A* and 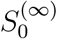 are constants. The first condition requires that *C*_0_ and *H*_0_ have the same specific growth rates, i.e.,

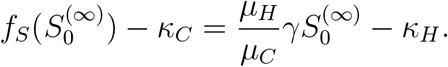

This equation has a unique solution for 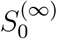 (assuming the condition that 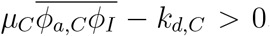, required for cyanobacterial viability) since *f*_*S*_(0) = 1 and *f*_*S*_(*S*_0_) is monotonically decreasing to 0 for *S*_0_ *>* 0. Then, given 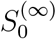, equation (16) requires, asymptotically, that

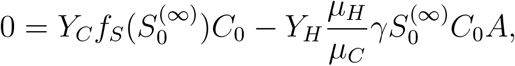

so that species ratio *A* is determined to be

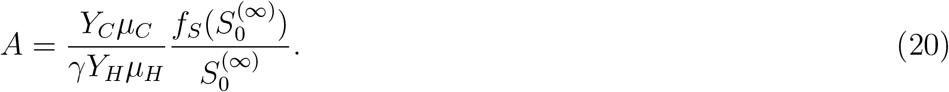

Relation (20) indicates that *A* is given by the ratio of cyanobacteria per capita production of byproduct to heterotroph per capita consumption of byproduct, taken asymptotically in the long-time system. For example, to maintain steady state, a relative increase/decrease in cyanobacteria per capita byproduct production would require a relative increase/decrease in heterotroph population. This can be counter-intuitive in some ways: for example larger *γ*, i.e., better environmental acclimation by heterotrophs than cyanobacteria, means smaller relative heterotroph population lest byproduct be depleted. However, though, more efficient use of byproduct by heterotrophs, i.e., smaller *Y*_*C*_, results in a larger relative heterotroph population.

#### 4.3.1 Computations

See Figure 9 for an example computational solution to equations (15)-(17), under non-viable conditions (top), viable conditions for cyanobacteria only (middle), and viable conditions for cyanobacteria and, eventually, for heterotroph invasion (bottom). Numerical simulations were performed using the MATLAB solver ‘ode15s’. See Table 2 for values of computational parameters.

**Table (2).**
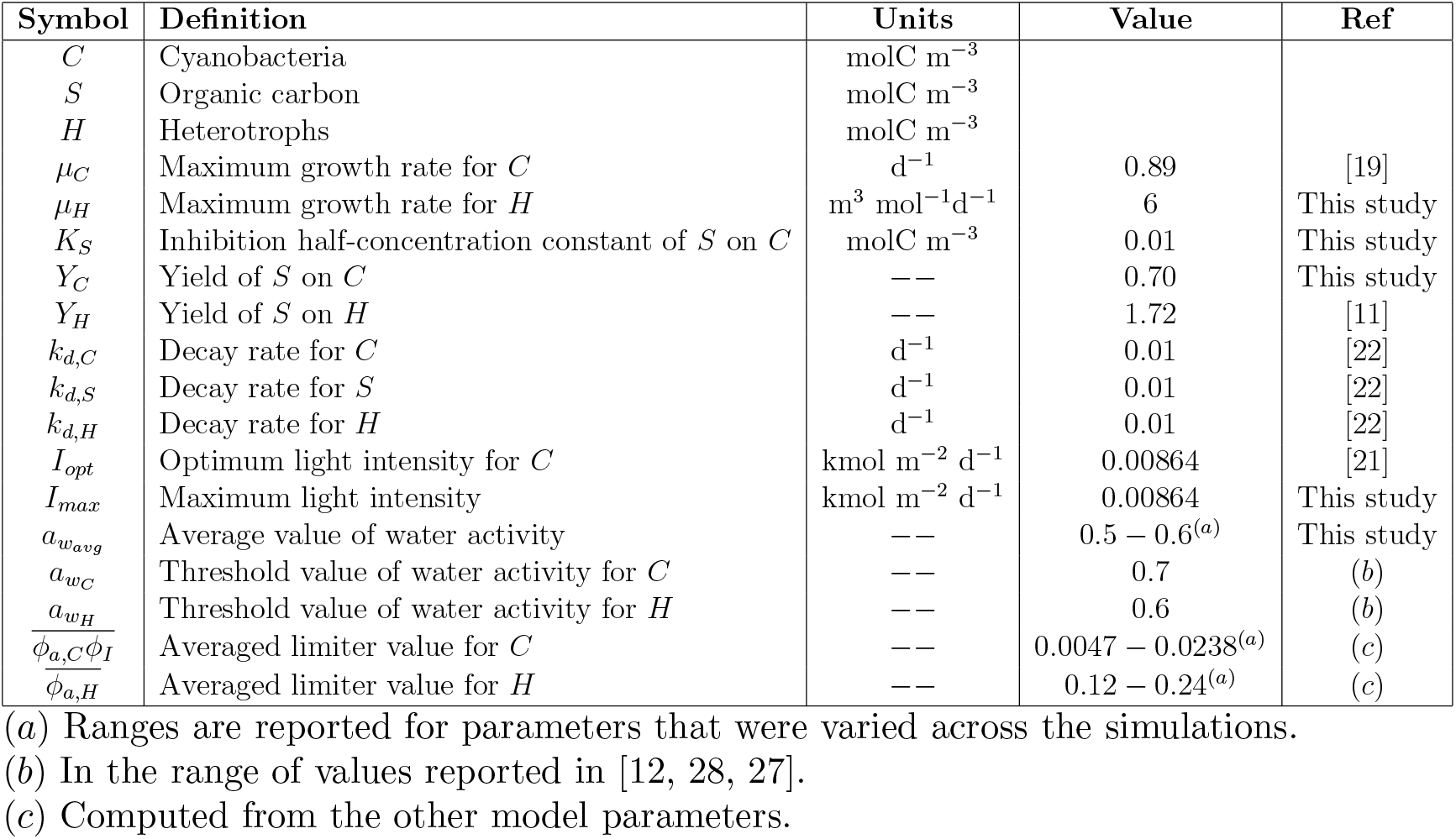
Variables and Parameters.

We make three principal observations in support of the previous discussion. First, even in the fully viable scenario, an established population of cyanobacteria is required before heterotrophs can successfully invade. This is a consequence of the need of heterotrophs for cyanobacterial byproduct, and, to an extent, the fact that byproduct does not quickly wash out of the system (here represented by a relatively small value of scaled decay coefficient *κ*_*S*_). Hence, observation of “young” cyanobacterial-only colonies is possible.

However, presence of a successful heterotroph population increases cyanobacterial biomass. This is not unexpected, as byproduct inhibition of cyanobacterial growth was explicitly included in the model. In fact, this inhibition is the only limiter (in the model) of cyanobacterial growth under viable conditions, so that introduction of heterotrophs is particularly dramatic: without a viable heterotroph population, cyanobacterial biomass is bounded, whereas as with heterotrophs, the cyanobacterial as well as heterotroph biomasses are unbounded. Obviously, something limits growth, but since we are not certain from available evidence what that something should be, we do not try to include an extra limiting mechanism. With regards to byproduct inhibition, direct evidence is scant [27] as *in situ* measurements are not currently available, but we consider the possibility reasonable as non-volatile compounds cannot easily escape SABs in many instances. We remark in any case that byproduct inhibition is not necessary to results reported here except as a mechanism by which cyanobacterial biomass can increase in the presence of heterotrophs.

Thirdly, certain model predictions are supported by observations reported here, including the presence *in situ* of both cyanobacterial as well as mixed cyanobacterial-heterotroph communities, e.g., see Figure 1. More notably, fixed ratios of cyanobacteria to heterotrophs are also observed, both as model predictions and from field measurements, recall Section 2. This, we propose, is a consequence of stoichiometric constraints, and, as far as we are aware, has been neither previously observed or predicted for SABs in the literature.

## 5 Discussion

Phototrophic SABs are commonly observed in light-exposed subaerial environments, including but not limited to stone, cement and other built material, e.g. [5, 6], though their broader roles in the surrounding and global ecosystems are not well understood. At the same time, SABs are interesting communities in and of themselves, apparently thriving under harsh exposed conditions where other biota cannot persist more than transiently.

A natural goal, then, is characterization of SAB community ecology and basic metabolic activity. How do SABs process the resources available to them? Though founded on cyanobacterial autophototrophic activity, these communities are also commonly observed to have heterotrophic inhabitants. Results of field studies at the Lincoln Memorial, Merchants’ Exchange Building, and the Federal Hall National Memorial also demon-strate the presence of mixed cyanobacterial-heterotroph communities, and have also, through microscopy at the micro-colony level, indicated a small proportion of cyanobacteria-only communities in addition to the more abundant mixed cyanobacteria-heterotroph ones. Interestingly, and further, the two sites have shown, backed by sequencing data, consistent ratios of heterotrophs to cyanobacteria in those mixed communities, even across the geographic range of the study sites.

The harsh environment of the typical SAB, in terms of aridity, temperature variation, UV exposure, etc., is not a focus here except through its oligotrophy, but could also be a fundamental factor in SAB ecology. Indeed, harsh environments have been suggested to support positive interactions between inhabitants [1], the so-called stress gradient hypothesis. For example, cyanobacteria could improve local conditions via water-attracting matrix structures or via pigments which reduce UV exposure or alter temperature in some way. However we see something apparently stronger than that: observations of cyanobacterial and heterotrophic biovolume, and supporting sequencing results, indicate similar SAB heterotroph/cyanobacteria ratios in different samples, even across three geographically separated sites. A simple model explains these observations in part, based on a supposition of overflow of photosynthetically fixed carbon. Such an overflow might arise for example if significant energy (from photosynthesis) is required for maintenance purposes due to extreme environmental stresses, resulting in a surfeit of fixed carbon. A similar overflow effect has been observed in cyanobacteria in a laboratory setting [3]; here we see evidence, albeit indirect, of it occurring in a natural population.

We go a bit further, based on laboratory studies that have in fact suggested that heterotrophs can result in increased population sizes, to propose that byproduct build up can lead to inhibition, for example through pH effects [22]. Supposing (in the model) that organic byproduct lowers pH and inhibits cyanobacterial growth, then the mathematical model also reproduces this ecological effect.

The model predicts that the heterotroph-to-cyanobacteria volume fraction is governed by per volume rates of production of excess carbon (by cyanobacteria) relative to per volume usage of excess carbon (by heterotrophs). The fact that we observe an approximately similar constant ratio, not only within individual study sites, but across all of our sites, suggests that there may be more to be understood. We cannot rule out some different coupling mechanism between the two populations, and we do not know to what extent this observation generalizes to phototrophic SABs beyond the ones studied here.

## 6 Acknowledgements

The authors would like to acknowledge the assistance of the US National Park Service as well as that of Madison Shoraka. Work presented here was funded through NSF/DMS grant 1951532, and through the auspices of the G.N.F.M. of I.N.d.A.M.

